# Signaling network model of cardiomyocyte morphological changes in familial cardiomyopathy

**DOI:** 10.1101/2021.08.28.458032

**Authors:** Ali Khalilimeybodi, Muhammad Riaz, Stuart G. Campbell, Jeffrey H. Omens, Andrew D. McCulloch, Yibing Qyang, Jeffrey J. Saucerman

## Abstract

Familial cardiomyopathy is a precursor of heart failure and sudden cardiac death. Over the past several decades, researchers have discovered numerous gene mutations primarily in sarcomeric and cytoskeletal proteins causing two different disease phenotypes: hypertrophic (HCM) and dilated (DCM) cardiomyopathies. However, molecular mechanisms linking genotype to phenotype remain unclear. Here, we employ a systems approach by integrating experimental findings from preclinical studies (e.g., murine data) into a cohesive signaling network to scrutinize genotype to phenotype mechanisms. We developed an HCM/DCM signaling network model utilizing a logic-based differential equations approach and evaluated model performance in predicting experimental data from four contexts (HCM, DCM, pressure overload, and volume overload). The model has an overall prediction accuracy of 83.8%, with higher accuracy in the HCM context (90%) than DCM (75%). Global sensitivity analysis identifies key signaling reactions, with calcium-mediated myofilament force development and calcium-calmodulin kinase signaling ranking the highest. A structural revision analysis indicates potential missing interactions that primarily control calcium regulatory proteins, increasing model prediction accuracy. Combination pharmacotherapy analysis suggests that downregulation of signaling components such as calcium, titin and its associated proteins, growth factor receptors, ERK1/2, and PI3K-AKT could inhibit myocyte growth in HCM. In experiments with patient-specific iPSC-derived cardiomyocytes (MLP-W4R;MYH7-R723C iPSC-CMs), combined inhibition of ERK1/2 and PI3K-AKT rescued the HCM phenotype, as predicted by the model. In DCM, PI3K-AKT-NFAT downregulation combined with upregulation of Ras/ERK1/2 or titin or Gq protein could ameliorate cardiomyocyte morphology. The model results suggest that HCM mutations that increase active force through elevated calcium sensitivity could increase ERK activity and decrease eccentricity through parallel growth factors, Gq-mediated, and titin pathways. Moreover, the model simulated the influence of existing medications on cardiac growth in HCM and DCM contexts. This HCM/DCM signaling model demonstrates utility in investigating genotype to phenotype mechanisms in familial cardiomyopathy.

## 1. Introduction

Familial cardiomyopathy, a common cause of heart failure (HF) and sudden cardiac death^1,2^, is a class of genetic disorder typically caused by mutations primarily in myocytes^3^. Dilated (DCM) and hypertrophic (HCM) cardiomyopathies are two main clinical phenotypes of familial cardiomyopathy, each affecting roughly from 1:500 to 1:2500 individuals globally^4–6^. To date, apart from transplantation, there are no effective cures for patients suffering from these cardiac muscle diseases^7^. Morphologically, DCM is characterized by dilation of the left ventricular chamber with systolic dysfunction. In contrast, HCM is associated with asymmetric cardiac growth, thickened ventricular walls, and diastolic dysfunction.

Even though numerous gene mutations that underlie HCM and DCM have been discovered^8^, our understanding of mechanisms leading from genotype to phenotype remains incomplete. This lack of understanding may originate from a significant diversity in mutations causing the same phenotype, especially in DCM^6^, or in the dependency of the cardiac phenotype on both environmental factors and genetic mutations^9,10^. However, the field has lacked systems approaches that integrate the relevant molecular signaling findings into a cohesive framework to scrutinize genotype to phenotype mechanisms. As cardiac signaling plays a crucial role in developing DCM and HCM^11^, understanding HCM/DCM signaling can elucidate major molecular players contributing to intrinsic variability and complexity of genotype-phenotype relations in familial cardiomyopathy. It may also help identify new drug targets and design more efficacious long-term pharmacological treatments for familial cardiomyopathy.

Although mutations in more than 100 genes have been linked to familial cardiomyopathy, sarcomeric gene mutations are the most common cause^12^. These mutations usually alter intracellular calcium dynamics, myofilament force generation, or both. In many cases, change in myofilament Ca^2+^-sensitivity is a specific alteration leading to contractile dysregulation. Often, DCM-associated mutations decrease myofilament sensitivity to calcium and lead to faster calcium dissociation from sarcomeres during relaxation and lower active force generation. In contrast, most HCM-linked mutations increase Ca^2+^-sensitivity followed by extended force relaxation time and higher contractile force^11,13^. Altered calcium handling and active force subsequently affect associated cardiac signaling pathways resulting in cardiac growth and remodeling.

In a significant study by Davis et al.^11^, the authors utilized a series of mouse models, *in vitro* and *in vivo*, combined with a computational model of myocyte contraction to study familial cardiomyopathy. They revealed that integrated active force magnitude over time is a significant predictor of cardiac phenotype during hypertrophic and dilated cardiomyopathies. They also showed that calcineurin (CaN)/nuclear factor of activated T cells (NFAT) and Ras/extracellular signal-regulated kinase 1/2 (ERK1/2) pathways are involved in developing HCM and DCM phenotypes by altering cardiomyocytes’ mass and eccentricity. An increase in the activity of other signaling components such as protein kinase B (AKT)^14^ and protein kinase G (PKG)^15^ have also been reported in experimental models of familial cardiomyopathy. Moreover, titin and its associated proteins and regulators such as FHL1, FHL2, and RBM20 act like a mechanosensor in cardiomyocytes and play a significant role in initiating and developing HCM and DCM through regulation of both molecular pathways and cardiac passive stiffness. While experimental models are necessary to identify mechanisms linking the genotype to phenotype in familial cardiomyopathy, challenges such as complex cardiac signaling pathways^16^, modified biomechanical stresses^14^, and neurohormonal system activity^17^ make them insufficient to determine the underlying mechanisms in HCM and DCM fully.

Computational models have been used to improve our understanding of mechanisms involved in multiple scales of cardiomyopathy^18^. However, at the cellular and tissue levels, while numerous signaling and finite element models have been developed for cardiac growth and remodeling^19^, to the authors’ knowledge, none of them investigated familial cardiomyopathies^18^. Hence, we constructed and validated a computational model of familial cardiomyopathy signaling on a cellular scale in this study. The model links the mutation-induced variations of calcium sensitivity to alteration of cardiomyocyte mass and eccentricity in hypertrophic and dilated cardiomyopathies. The model focuses on signaling related to differences in cardiomyocyte remodeling between HCM and DCM rather than other clinical features such as systolic and diastolic function, fibrosis, apoptosis or myofiber disarray due to multi-scale systems complexity and limited data availability. The model indicates the significant signaling reactions regulating the cardiomyocyte morphology in HCM/DCM mutations altering active force though calcium sensitivity and provides new insights on potential drug targets to decrease cardiomyocytes’ adverse growth and remodeling in familial cardiomyopathy.

## 2. Results

### 2.1 A conceptual model of cardiac remodeling in response to HCM/DCM mutations and mechanical loading

In terms of cell morphology, the main difference between HCM and DCM is cardiomyocyte eccentricity. HCM mutations cause cardiomyocyte growth through the addition of sarcomeres in parallel that increases myocyte cross-sectional area without significant changes in length. On the contrary, DCM mutations increase myocyte length by adding sarcomeres in series. Hence, HCM and DCM mutations decrease and increase myocyte eccentricity, respectively. Among the primary factors of familial cardiomyopathy regulating myocyte eccentricity, several studies demonstrated the integrated active force as a key factor^11,20^. HCM and DCM affect cardiomyocyte active force through changing myocyte contractility^21^. The negative correlation between myocardial active force and eccentricity is not limited to familial cardiomyopathy. In the pressure overload condition, elevated afterload increases active force^22^ and decreases myocardial eccentricity^23^ (Fig. 1a).

**Fig. 1.**
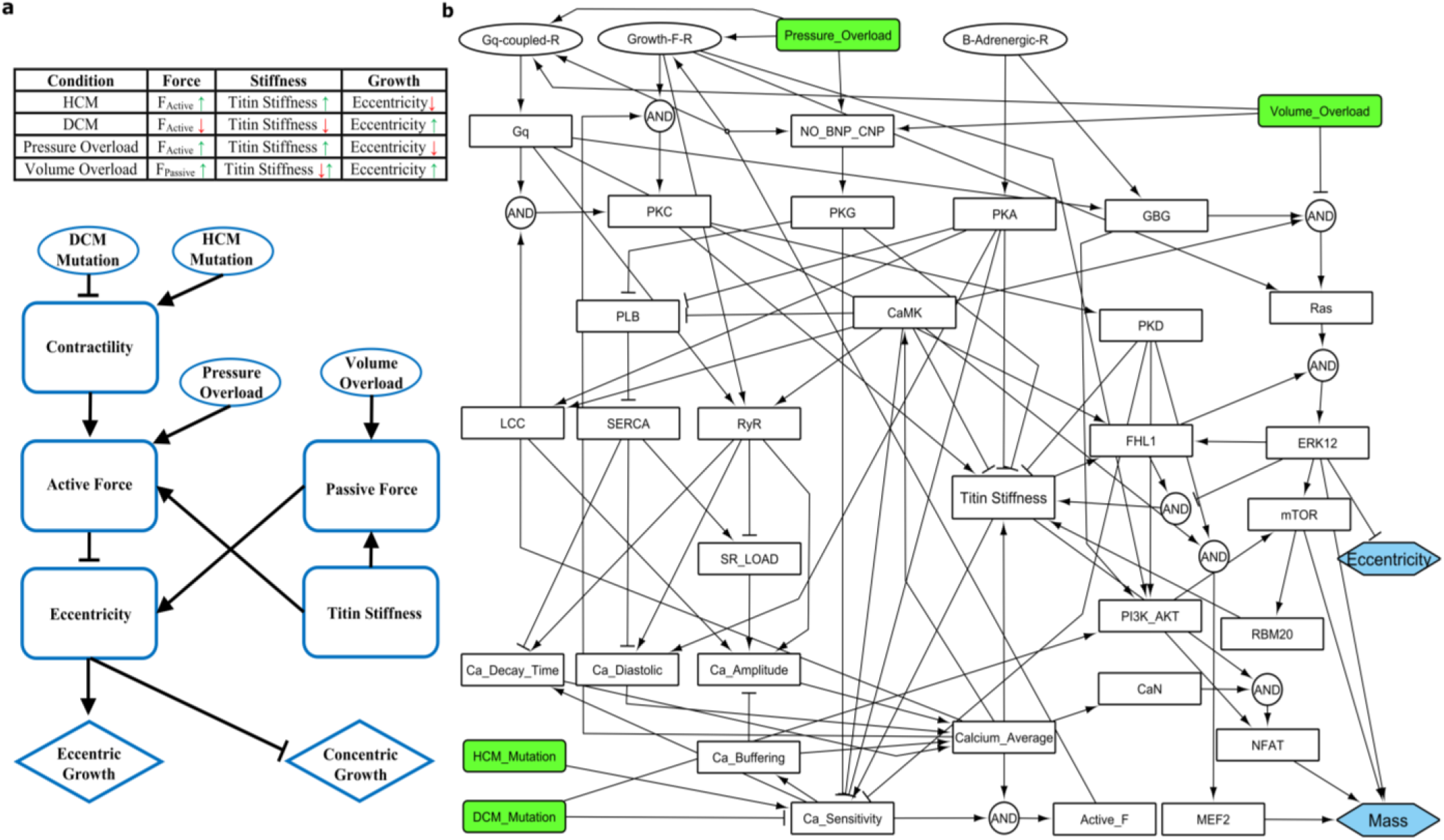
Reconstruction of the familial cardiomyopathy signaling network. (a) Context-dependent responses of cardiomyocytes in cardiac growth and remodeling are simplified by a conceptual model of HCM/DCM. Interactions between cardiomyocyte active force, passive force, titin stiffness, and eccentricity regulate cardiac phenotype. (b) HCM/DCM signaling network. Four model inputs (green) are linked to two model outputs in the myocyte (blue) through 33 signaling species (white) and 82 reactions.

In the context of volume-overload, increased cardiac passive force regulates myocyte eccentricity. After volume overload, there is a shift of the end-diastolic volume to the right in the ventricular pressure-volume loop, which results in higher cardiac tissue elasticity and passive force. This increased passive force then initiates a cardiac response leading to cardiac dilation^23^. Interestingly, titin stiffness is a significant contributor to cardiomyocyte passive stiffness and force and is positively correlated with active force in HCM, DCM, and pressure-overload conditions. Yet, titin stiffness is negatively correlated with eccentricity except in some reported cases in volume-overload hypertrophy (Fig. 1a). These observations suggest a potential mediating role for titin stiffness in genotype to phenotype mechanisms. We propose a conceptual model (Fig. 1a) that can explain the distinct cardiac responses observed in various contexts by considering the above factors and their interrelations. The model emphasizes the role of titin and biomechanical factors such as active and passive forces in regulating familial cardiomyopathies. However, in cardiomyocytes, a complex signaling network conveys the effects of gene mutations on cell eccentricity and stiffness are and is responsible for interactions illustrated in the conceptual model.

### 2.2 Predictive computational model of HCM/DCM signaling network

To reconstruct the familial cardiomyopathy signaling network (Fig. 1b), we identified the major signaling modules contributing to cardiomyocyte responses in familial cardiomyopathy and reconstructed the molecular interactions between signaling components by employing the cardiac signaling networks developed by Ryall et al.^24^ and Tan et al.,^25^ which investigated cardiac hypertrophy and mechano-signaling, respectively. Due to limited data availability, we included components with demonstrated roles in cardiac growth and remodeling and available experimental data or perturbations in studied contexts (Fig. 1b). Regarding the main signaling modules, Davis et al. ^11^indicated the myofilament force-generation module, CaN-NFAT, and ERK1/2 pathways regulate mass and eccentricity in familial cardiomyopathy. In addition, other signaling pathways such as PI3K-AKT^14,21^, NO-PKG^15,26^, calcium^27^, β-adrenergic^13,28^, and downstream pathways of Gq protein^29^ and growth factor receptors^30,31^ could play a role in familial cardiomyopathy. Moreover, titin has been described as a regulator of familial cardiomyopathy via its effects on cardiomyocyte stiffness, force generation^32^, and mechanotransduction^33^.

Since titin and calcium modules were less-developed in previous models^24,25^, we reconstructed them with more cell signaling detail. As shown in previous studies^34^ and proposed by the conceptual model (Fig. 1a), titin could play a major role in regulating cardiac signaling in familial cardiomyopathy. Several kinases like Ca^2+^/calmodulin-dependent protein kinase (CaMK), protein kinase D (PKD), protein kinase A (PKA), ERK1/2 phosphorylate titin at different positions and decrease titin stiffness. In contrast, phosphorylation by protein kinase C (PKC) increases titin stiffness^26^. In addition, RNA binding motif protein 20 (RBM20) regulates alternative splicing of titin, which changes the ratio of compliant N2BA to stiff N2B isoform and subsequently affects titin stiffness. Titin also can affect ERK1/2 and CaN-NFAT pathways through its associated proteins, including four and a half LIM domain (FHL) proteins, FHL1 and FHL2^35^. The intracellular calcium cycling module is a significant part of active force generation in cardiomyocytes, and its diverse response has been reported in familial cardiomyopathy^11,27^. Also, mutations in calcium regulating proteins such as phospholamban (PLB) might lead to familial cardiomyopathy^36^. Therefore, we added a simplified calcium module comprising its related characteristics such as sarcoplasmic reticulum (SR) Ca^2+^ load, Ca^2+^ amplitude, buffering, diastolic level, and decay time, as well as its regulating proteins and channels such as PLB, SR Ca^2+^-ATPase (SERCA), ryanodine receptors (RyR), and L-type calcium channel (LTCC). It is noteworthy that we added pressure overload and volume overload stimuli to our model to explore the dynamics of familial cardiomyopathy signaling influenced by biomechanical signals. This enables the model to simulate the influence of some medications on cardiac phenotype, capturing both their molecular and hemodynamic effects.

Overall, figure 1 shows the signaling network model developed in this study. This model links the mutation-induced changes in calcium sensitivity to variations in cardiac signaling pathways and shows how these variations result in changes in mass and eccentricity of cardiomyocytes. The HCM/DCM signaling network model includes 39 nodes (four inputs: HCM mutation, DCM mutation, pressure overload, volume overload; two outputs: mass and eccentricity; and 33 signaling species) connected by 81 reactions (S1 Table). Applying a logic-based differential equations (LDE) approach^37^, we converted the HCM/DCM signaling interaction graph (Fig. 1b) into a predictive computational model. The normalized activity of each node is computed utilizing ordinary differential equations with saturating Hill functions. We used continuous logical “AND” and “OR” gates to combine the influence of multiple reactions on a signaling node. Similar to our previously published models of cardiac signaling^16,24^, constant default values have been used for all network parameters. To prevent saturation in unstimulated conditions, we used a smaller default value for reaction weight of inhibitory reactions reducing calcium sensitivity and titin stiffness. Based on the availability of quantitative experimental data, individual parameters of the model can be calibrated subsequently to provide more quantitative predictions. Using Netflux software, we automatically generated a system of LDEs and simulated HCM/DCM signaling dynamics in MATLAB.

### 2.3 Model performance in predicting context-specific qualitative data

To evaluate the model performance in predicting cardiac morphology and variations of signaling nodes in four contexts (hypertrophic and dilated cardiomyopathies, pressure-overload, and volume-overload hypertrophy), we curated context-specific experimental data from the literature. All data were categorized as a decrease, no change, and increase according to their statistical significance compared to the control in each study (S2 Table). As diversity in test data is crucial for assessing the performance of complex models, we employed the following strategies in obtaining data from the literature. First, we tried to obtain data, as much as possible, for all key signaling nodes of the model in studied contexts such as HCM and DCM. Second, various experimental data with and without intermediate perturbations such as inhibition, overexpression, and multi-stimuli were obtained. Due to their importance, we also included overexpression data (no additional context) and Gq context data in the validation dataset. Third, we had all reported responses for signaling nodes in a context, such as both calcium amplitude’s increase and decrease in DCM (Fig. 2).

**Fig. 2.**
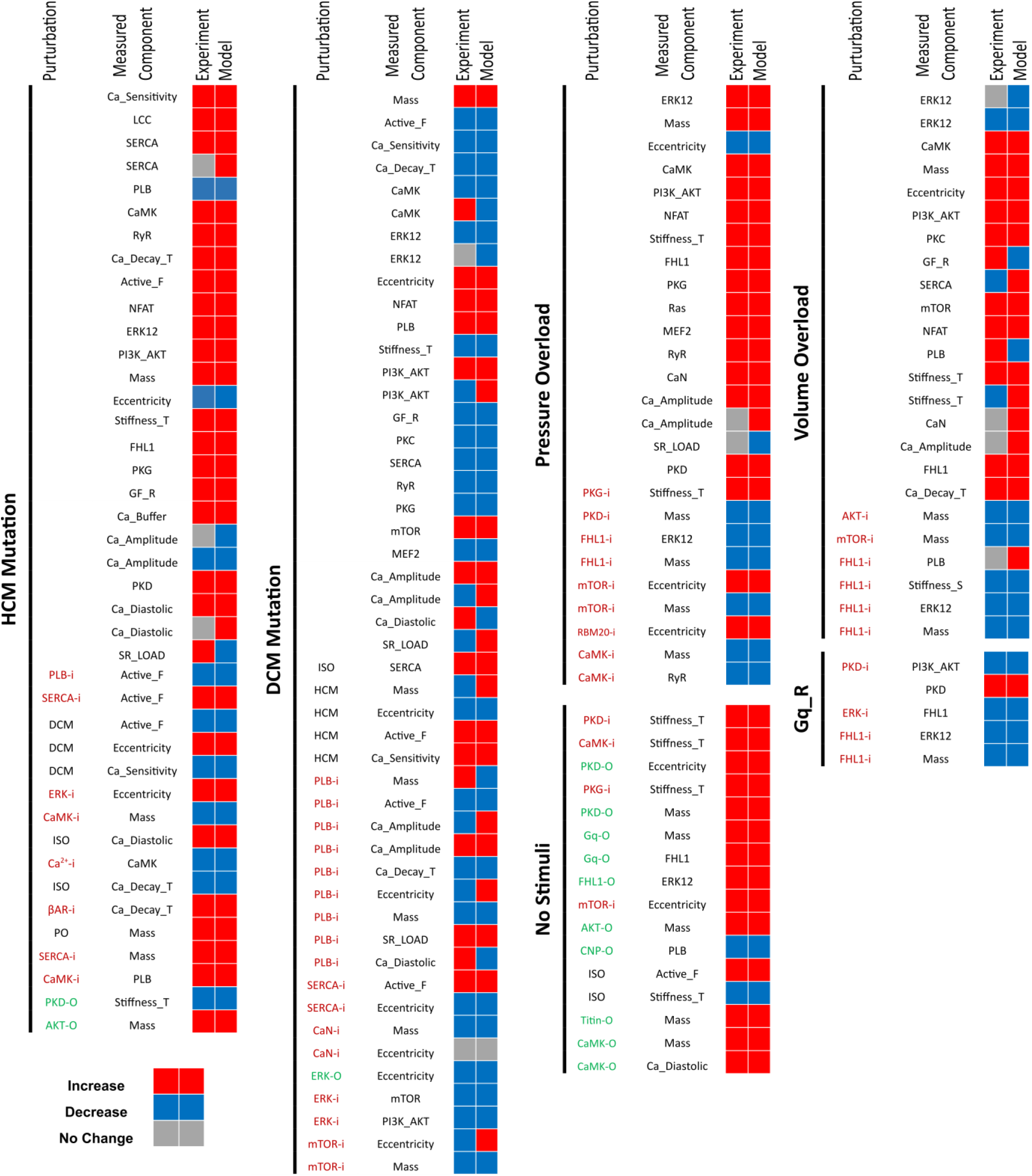
Classified qualitative validation illustrates HCM/DCM model performance in different contexts. Comparison of model predictions with 160 experimental data points from literature (S2 table). Inhibitory and overexpression perturbations are illustrated by −i (red) and −o (green), respectively. For model results, relative changes that are higher than the *in-silico* threshold (0.001%), and for experimental data, statistically significant changes compared to the control have been considered an increase/decrease.

Simulated activity changes of network nodes after HCM mutation, DCM mutation, pressure overload, and volume overload with/without perturbation of various nodes are compared with associated validation data in Fig. 2. The HCM/DCM model with default parameters has a total prediction accuracy of 83.8% (134/160) across all contexts. The model has its highest prediction accuracy in the pressure overload context with 92.3% (24/26), followed by HCM with 90.2% (37/41), DCM with 75% (36/48), and the volume overload context with 66.6% (16/24) validation accuracy. Prediction accuracy for each context was determined by dividing the number of model’s correct predictions by the number of all experimental data points in each context (see Fig. 2).

It is noteworthy that discordant responses reported for the same experiment decrease the model prediction accuracy regardless of actual model performance. These nodes are mostly located in the calcium module. For example, while Davis et al.^11^ reported increased Ca^2+^ transient amplitude in DCM, Robinson et al.^21^ and Liu et al.^38^ reported the opposite. This inconsistency could result from different gene mutations^6^ or altered mechanisms regulating calcium level^39^. In the case of discordant experimental data, all data were included as separate validation tests. The DCM context has the highest number of conflicting experimental data. In addition to Ca^2+^ transient amplitude, CaMK, ERK1/2, and PI3K-AKT responses to DCM mutations are also contradicting in the literature. However, changes in mass and eccentricity as the main indicators of the cardiac phenotype are consistent in each context and correctly predicted by the model.

As supported by model results and experimental observations (Fig. 2), HCM mutations increase calcium sensitivity, integrated active force, CaMK, PKG, PI3K-AKT, ERK1/2, NFAT, and titin stiffness (S2 Table). HCM mutation consequently leads to an increase in mass and a decrease in eccentricity. In DCM mutation, the model shows opposite changes in CaMK, PKG, ERK1/2, and titin stiffness, but there are still increases in PI3K-AKT, NFAT, and mass. In addition, the model indicates attenuated changes in eccentricity after concurrent HCM and DCM input signals. Powers et al.^20^ and Davis et al.^11^ reported that double-transgenic murine models containing HCM and DCM gene mutations have more normal cardiac morphology than DCM or HCM murine models. Regarding titin stiffness, the model simulation results illustrate an increase in titin stiffness after HCM mutation, pressure overload, and volume overload, as shown by Herwig et al.^29^, Michel et al.^40^, and Mohammad et al.^41^, respectively. In contrast, DCM mutations lead to a decrease in titin stiffness^42^.

### 2.4 Identification of key network regulators and potential interactions in cardiac remodeling

A key step in finding effective drugs for a disease is determining how alteration in individual reactions will affect the cell response. Determination of reactions with the highest impact reveals the significance of each pathway in regulating the cell response and minimizes the efforts and costs of follow-up experiments by avoiding non-significant pathways. Although local sensitivity analysis can provide useful information^43^, it does not capture nonlinear interactions between reactions in a complex network like HCM/DCM signaling. Thus, we performed Morris global sensitivity analysis by applying a change in a reaction weight and calculating its global effect on the network to identify reactions with the highest impact on model prediction accuracy, as shown in Fig. 3. By selecting the reactions’ weight (W) as the variable parameter, we calculated the Morris index (μ*) and standard deviation for each reaction in the model. Reactions with higher scores (μ*>2) were visualized on the HCM/DCM signaling network (Fig. 3). As illustrated in Fig. 3, 24 of 82 model reactions significantly affect the model prediction accuracy.

**Fig. 3.**
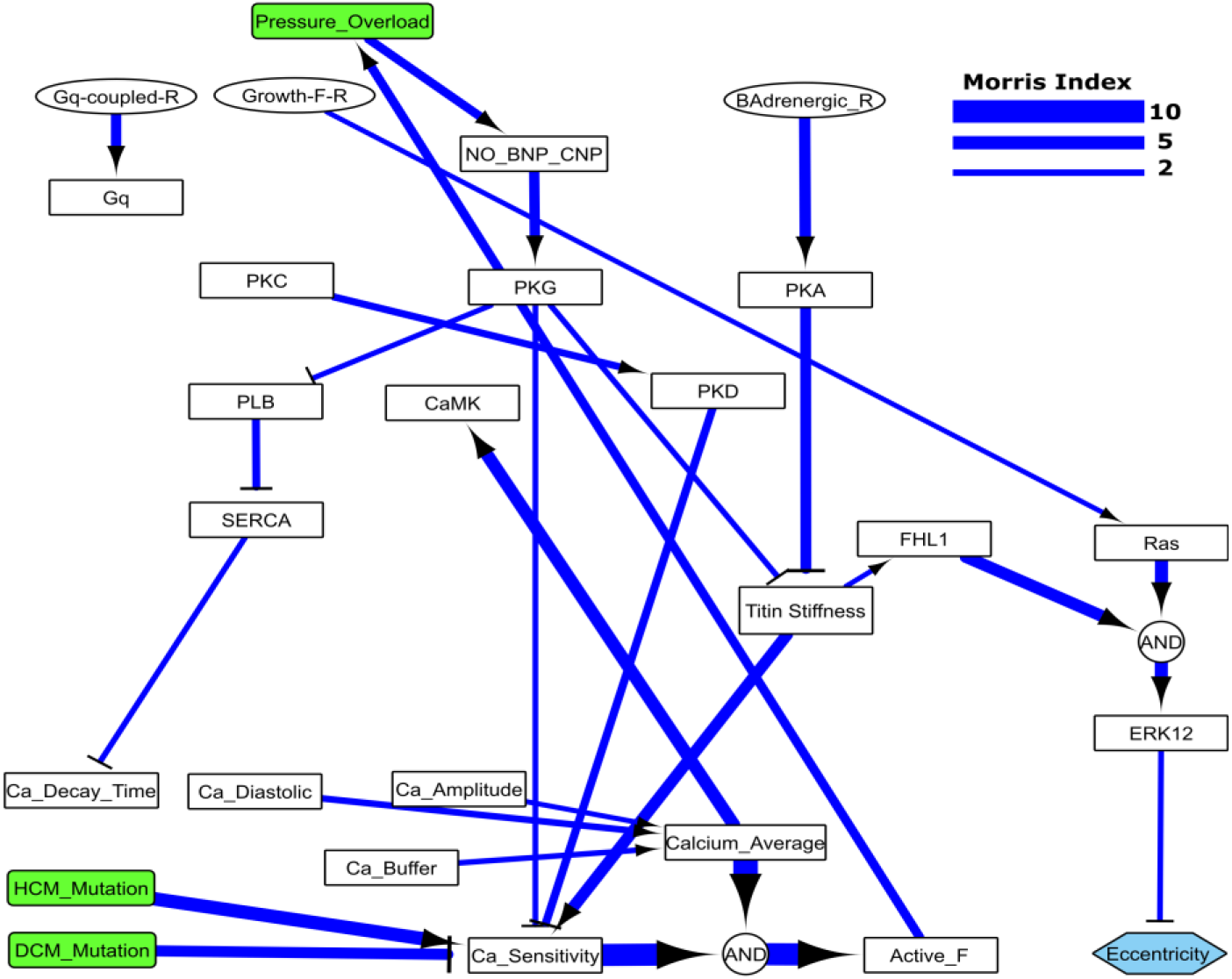
Morris global sensitivity analysis identifies key signaling reactions in familial cardiomyopathy. (a) The Morris index (μ*) depicts important (μ* >2) and less-important reactions in HCM/DCM signaling model. Greater Morris index means more influence on model validation percent. A larger σ to μ* ratio demonstrates a more nonlinear effect. (b) Visualization of important reactions in familial cardiomyopathy signaling based on their ranks from the Morris sensitivity analysis. The thicker arrows illustrate the larger Morris index.

Although non-significant reactions may partly contribute to signal transduction in cardiomyopathy, modifying them has a negligible effect on the overall response of the HCM/DCM signaling network. For example, CaMK, PKA, PKG, ERK1/2, and PKC phosphorylate titin and modify its stiffness. However, based on the results, only PKA and PKG effects on titin stiffness have significant contributions to HCM/DCM signaling. It is noteworthy that significance in the Morris sensitivity analysis is a relative concept. Based on the sensitivity analysis from Fig. 3, reactions modeling the effects of calcium sensitivity and calcium average on the active force have the highest impact on HCM/DCM signaling network. Powers et al.^20^ and Davis et al.^11^ also reported significant changes in cardiac phenotype after a modification in calcium sensitivity and active force in HCM or DCM contexts. Moreover, as model results indicate, activation of CaMK by calcium^27^, titin contribution to active force through calcium sensitivity^32^, activation of PKA by β-adrenergic receptors^44^, PKG by nitric oxide^45^, and ERK1/2 by Ras and FHL1^46^ could significantly affect cardiac response in cardiomyopathy.

Even though the HCM/DCM signaling model has a reasonably good validation accuracy (83.8%), there are some incorrect predictions. Apart from discordant responses for the same experiment, imprecise model parameters and structure could influence model prediction accuracy. Because of limited quantitative data availability, we used default parameters for the model, which inevitably reduces the model prediction accuracy. Concerning model structure, we developed the model by employing identified mediators of cardiac signaling in familial cardiomyopathy with their known interactions in cardiac signaling. Thus, new yet-to-be-discovered interactions may exist in cardiomyopathy signaling. We applied a structural revision method, CLASSED, which we developed previously^16^ to find new potential interactions that improve model validation accuracy. We screened all possible interactions between the model components by a systematic, one-by-one adding of new reactions between signaling nodes through the “OR” gate and from each node to the existing reactions by an “AND” gate.

Figure 4 displays the inferred reactions that improve the model prediction accuracy by more than one percent. These inferred reactions indicate potential causal relations between signaling components and provide a baseline for further experiments. To obtain figure 4, we added new reactions to the model one by one to explore what reactions can increase the model validation accuracy. The reactions with the highest rise in model validation accuracy are potential reactions that may contribute to cardiomyocyte response but are not identified before by previous experimental studies on cardiomyocytes. According to Fig. 4, Gq protein and volume overload stimulus may affect calcium signaling by raising PLB activity. The HCM/DCM model (Fig. 1b) shows that PKC is a common downstream component of Gq protein and volume overload signals and may mediate these inferred reactions. Braz et al.^47^ showed that PKC affects PLB phosphorylation and activity. PLB phosphorylation increased in PKCα-deficient mice, whereas it decreased in transgenic mice overexpressing PKCα.

**Fig. 4.**
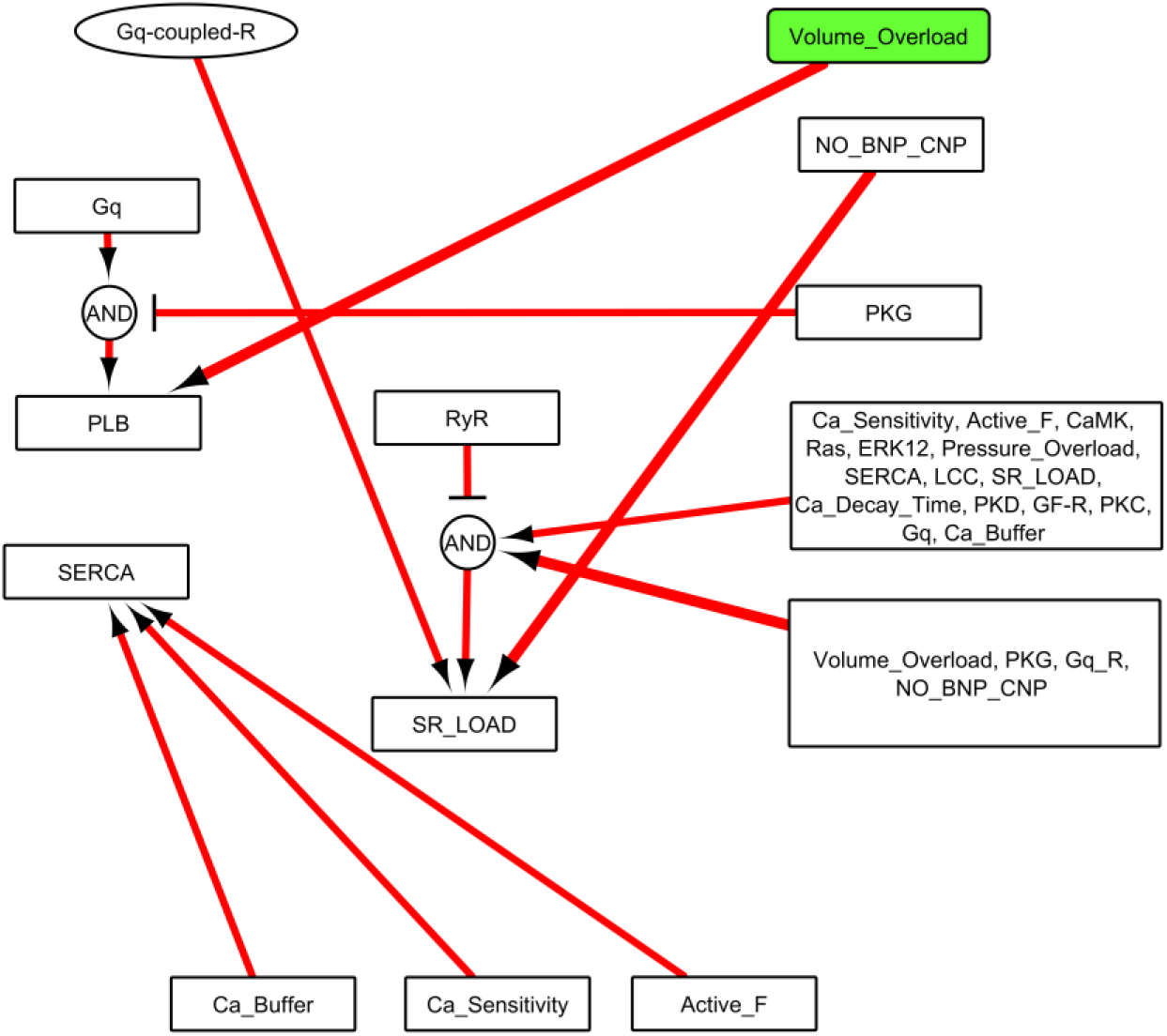
Structural revision analysis suggests new potential interactions in HCM/DCM signaling. The thicker arrows illustrate a larger improvement in model prediction accuracy.

The results also suggest contributions of several signaling nodes such as NO and PKG in SR Ca^2+^ load regulation (Fig. 4). In the model, NO leads to phosphorylation of PLB by PKG and increases Ca^2+^ uptake by SERCA, consequently raising SR Ca^2+^ load and Ca^2+^ transient amplitude. Thus, the inferred reactions point to a more pronounced positive effect of NO-PKG on SR Ca^2+^ load in familial cardiomyopathy through other parallel pathways and higher reaction weights (W) for the existing pathway. This agrees with preclinical studies. While both positive and negative effects of NO-PKG on SR Ca^2+^ load and Ca^2+^ transient through mediators like SERCA, RyR, sodium/hydrogen exchanger (NHE), PKA, PLB have been reported^48–50^, its positive effect seems to be dominant in cardiomyopathy signaling^51^. Previous studies reported increased activity of PKG and SERCA with elevated SR Ca^2+^ load in HCM and the opposite _response in DCM_^26,28,52^.

### 2.5 Evaluation of new and existing pharmacotherapies

A growing number of clinical trials on combination therapies^53^, and their higher performance in treating cardiac disease like heart failure^54^, make them an important consideration for drug development. Thus, we analyzed the efficiency of all possible drug targets in the context of HCM and DCM mutations through combinatory inhibition and overexpression of model signaling nodes by changing the Ymax parameter from 1.0 to 0.5 and 1.5, respectively.

As shown in Fig. 5a, drugs that knock down two targets have the highest effect in decreasing mass and increasing eccentricity in the HCM context. In DCM, combined pharmacotherapies including concurrent overexpression and knockdown of two separate nodes provide the best results in preventing cardiac growth and remodeling in DCM. For example, in the HCM context, paired knockdown of growth factor receptors (GF-R) with calcium, titin stiffness, or G beta-gamma complex (GBG) leads to the highest decrease in mass and increase in eccentricity. In general, combined knockdown of nodes such as GF-R, PI3K-AKT, NFAT, ERK1/2, mammalian target of rapamycin (mTOR), titin stiffness, and calcium are predicted as potential treatment strategies for the HCM context. In the DCM context (Fig. 5b), the number of potential drug targets is less than the HCM context, and usually, they are limited to knockdown of NFAT or PI3K-AKT with overexpression of Gq protein-associated receptors (Gq-R), calcium, ERK1/2, FHL1, and titin stiffness.

**Fig. 5.**
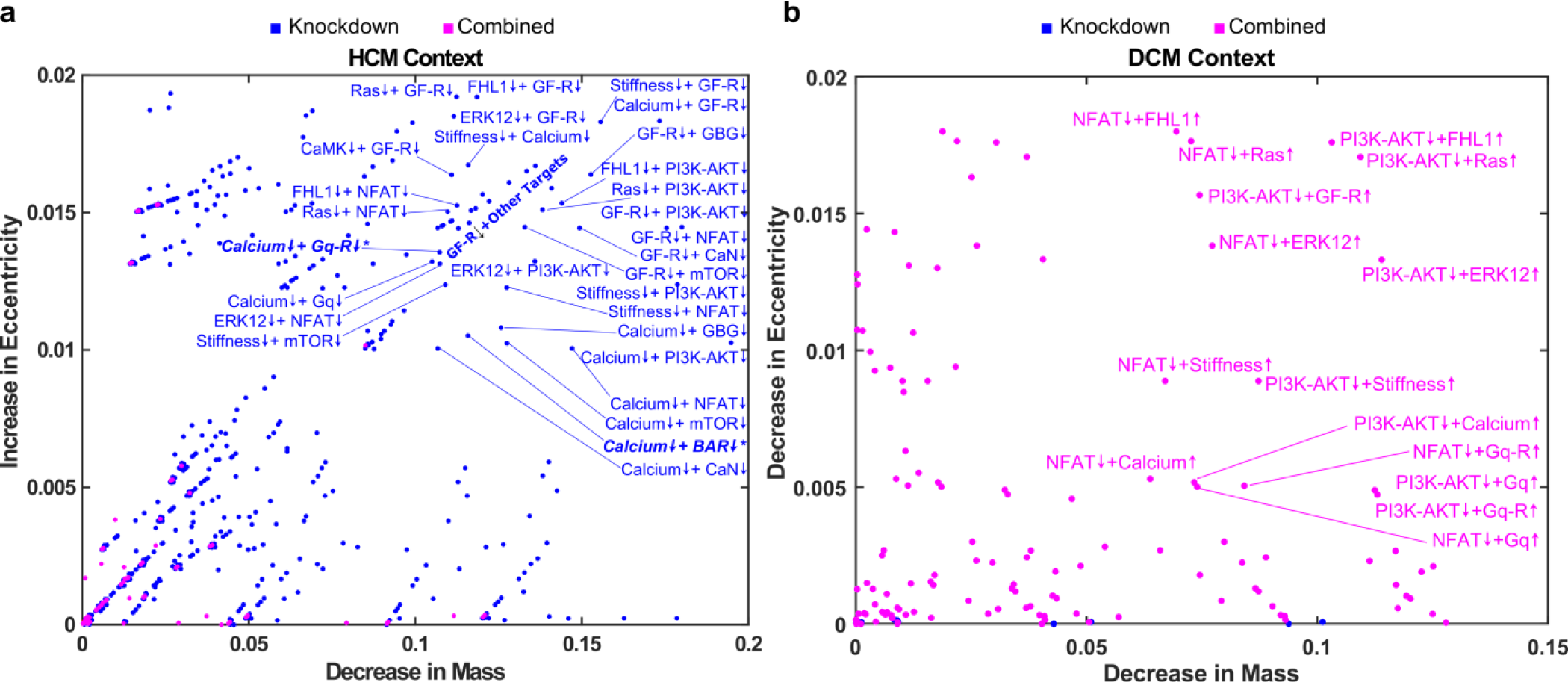
Efficacy of combination pharmacotherapies on familial cardiomyopathy. All pairwise combinations of reducing (Ymax=0.5) or increasing (Ymax=1.5) nodes’ activity and their subsequent effects on myocardial mass and eccentricity in the context of HCM mutations (a) and DCM mutations (b). (*) shows two combinatory drug targets (Calcium+Gq-R, Calcium+βAR) among conventional treatment strategies.

While no previous experimental studies investigated combination pharmacotherapies for familial DCM or HCM, some studies showed the effectiveness of targeting one of the predicted signaling nodes by the model. As shown in Fig. 5, the model predicts that ERK1/2 overexpression or inhibition can contribute to preventing mutation-induced changes in cell eccentricity in DCM and HCM, respectively. Likewise, Davis et al.^11^ observed that mitogen-activated protein kinase kinase (MEK) overexpression (upstream of ERK1/2) in rat cardiomyocytes decreases the DCM-induced increase in cell eccentricity. They also showed that MEK inhibition reverses the decrease in cell eccentricity in rat cardiomyocytes with HCM mutation.

In a recent study by Bu et al.^55^, the authors investigated seven cardiomyopathy-related signaling pathways and showed that pharmacological inhibition of mTOR, MEK-ERK, and β-adrenergic signaling and genetic inhibition of mTOR, MEK-ERK signaling could rescue HCM phenotypes in zebrafish model. Based on our model predictions, ERK inhibition resulted in an increase in eccentricity and a moderate decrease in mass, consistent with experimental observations. Moreover, the model predicted that mTOR inhibition results in a substantial decrease in mass and no significant increase in eccentricity also consistent with experiments. The model predicted β-AR inhibition may have a moderate effect on rescuing HCM phenotypes (mass and eccentricity). Experimental results from the zebrafish model with β-AR inhibition showed partial rescue of the HCM phenotype, but this effect was not significant with genetic inhibition of β-AR pathway. Based on model predictions, CaN inhibition may moderately reduce mass but would not affect eccentricity. Experimental results show some improvement in cardiac function when CaN is pharmacologically inhibited. Consistent with experimental results, the model also predicted minor or no significant changes in HCM phenotypes after CaMKII and HDAC-MEF2 inhibition^55^.

Furthermore, the model predicted two known targets of existing heart failure drugs. As shown in Fig. 5a, the combined knockdown of calcium with βAR or Gq-R could prevent the HCM phenotype. Ho et al.^56^ and Semsarian et al.^57^ showed in their studies that diltiazem, a Ca^2+^ channel blocker, has the potential to ameliorate the adverse remodeling in familial hypertrophic cardiomyopathy in pre-clinical and mouse model studies, respectively. However, it is important to consider the limitations of available pre-clinical studies in covering large groups of patients or diverse mutations^56^ or discordant results in murine models for the effectiveness of Ca^2+^ channel blocker in reversing cardiomyopathy phenotype^57,58^. Thus, the recent AHA/ACC 2020 guideline^59^ could not provide conclusive evidence to support the effect of calcium channel blockers in HCM.

While the model suggests several targets for combination pharmacotherapy of familial cardiomyopathy, it is important to consider the effects of modified targets on other aspects of cardiomyocyte response such as fibrosis, apoptosis, and fetal genes expressions before moving forward with experiments. For example, activation of Gq-receptor pathways suggested by the model as a part of a therapeutic approach in DCM influences many other molecular mechanisms in addition to cardiomyocyte growth. While the effect of Gq-receptor activation on increasing ERK activity and potential hypertrophic growth^60,61^ could be beneficial for DCM by reducing cardiomyocyte eccentricity, its adverse effects on cardiac fibrosis^62,63^ and apoptosis^60,64^ could contradict the favorable outcomes.

To validate a potential drug combination predicted by the model, we conducted an experiment on recently reported patient-specific iPSC-derived cardiomyocytes (iPSC-CMs) harboring two heterozygous missense mutations, one in β-myosin heavy chain 7 (MYH7-R723C), which expressed in sarcomere of cardiomyocytes and the other one in muscle LIM protein (MLP) expressed on Z-disc of sarcomeres^65^. This patient with double mutant (MLP-W4R;MYH7-R723C) was diagnosed with HCM phenotype at the age of 1.3 years^65^ and iPSC-derived CMs from this patient exhibited significantly elevated expression of HCM markers including *ANF* and *BNP* mRNA levels and enlarged cell area phenotype^65^. Among drug targets predicted by the model in the HCM context (Fig 5A), we selected combination inhibition of ERK1/2 and PI3K-AKT pathways for experimental validation for their efficacy as well as feasibility with existing compounds. 25 days old iPSC-CMs were treated with 0.5 μM MEK-ERK inhibitor (U0126-EtOH, IC_50_: 60-70 nM), 0.2 μM PI3K-AKT inhibitor (Wortmannin, IC_50_: 2-4 nM), or combination of both inhibitors. DMSO was used as vehicle control. The reversal of HCM phenotype was assessed by measuring expression levels of *ANF* and *BNP* mRNA after one day drug treatment as well as the cell area of iPSC-CMs after 4 days of drug treatment.

Fig. 6a shows iPSC-CMs after all four treatments. As illustrated in Figures 6b and 6c, no significant decrease was observed for *ANF* and *BNP* mRNA with single drug treatment by ERK or PI3K-AKT inhibitors. Interestingly, in line with our model prediction, combined inhibition of ERK+PI3K-AKT pathways resulted in a significant decrease in *ANF* and *BNP* mRNA expression levels. Moreover, the decline in cell area was also more pronounced in iPSC-CMs with a combined inhibition of ERK+PI3K-AKT (Fig. 6d) than the ERK inhibition individually and no significant decrease for PI3K-AKT inhibition. These data thus successfully validated the model prediction on combined inhibition of ERK+PI3K-AKT pathways as a potentially new rescue strategy for HCM.

**Fig. 6.**
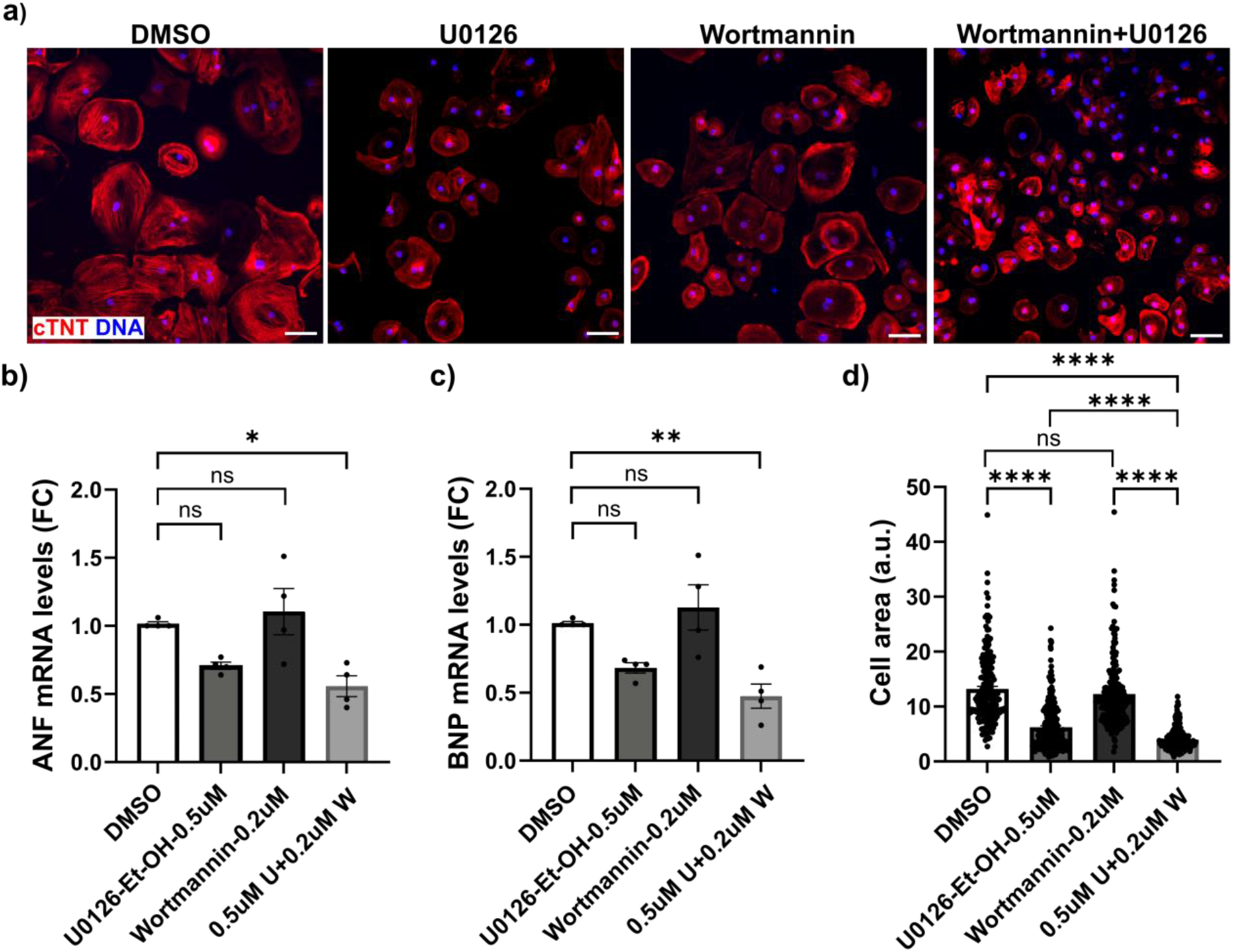
Reverse of HCM phenotype by Combined inhibition of ERK + PI3K-AKT pathways. Treatment of iPSC-cardiomyocytes (CMs) derived from iPSC of the double mutant (MLP-W4R;MYH7-R723C) HCM patient with MEK-ERK inhibitor and PI3K-AKT inhibitor. a) Immunostaining of cTnT (red; cardiac specific protein) in MLP-W4R; MYH7-R723C iPSC-CMs treated with DMSO (vehicle), 0.2 μM of Wortmannin, 0.5 μM of U0126-EtOH, and combined treatment of 0.2 μM of Wortmannin + 0.5 μM of U0126-EtOH on day 25 for four days (Scale bar, 50 μm). *ANF* (b) and *BNP* mRNA (c) levels significantly decreased in combined inhibition of ERK + PI3K-AKT pathways. ERK inhibition and combined inhibition of ERK + PI3K-AKT pathways reversed HCM mutation-induced changes in cell area (d) of iPSC-CMs. All data points derived from at least 3 independent cardiomyocyte differentiation batches. All data are presented as mean±SEM; *P<0.05; **P<0.01; ***P<0.001; ****P<0.0001.

Considering that iPSC-CMs on unpatterned substrates are not elongated as seen *in vivo*, they cannot be examined to determine changes in cell eccentricity. While we observed some changes in cell shape characteristics such as altered cell circularity for ERK inhibition and combined inhibition of ERK+PI3K-AKT, these changes are not directly comparable to the model predictions of eccentricity, which would require *in-vivo* experiments or experiments with iPSC-CMs seeded on predesigned scaffolds^65^.

Numerous drugs have been developed to relieve the symptoms of and prevent heart failure progression^67^. While some partly prevented cardiac growth and remodeling in heart failure^68^, a few studies focused on the effects of current drugs on cardiac remodeling in familial cardiomyopathies. Hence, we employed the HCM/DCM signaling model to explore the outcomes of available heart failure medications and two new categories of drugs (myosin inhibitors and activators) on cardiomyocyte morphology explained by changes in cardiomyocyte mass and eccentricity (Table 1). It is noteworthy that none of these drugs are approved by FDA for familial cardiomyopathy. We categorized drugs based on their mechanism of action and included hemodynamic effects to increase simulation accuracy. As shown in Table 1, the model simulations indicate a decrease in mass and an increase in eccentricity in HCM and DCM contexts for most medications. Even though a reduced mass is a positive outcome for patients with HCM and DCM, increased eccentricity would be detrimental for DCM patients. In contrast, cardiac glycosides such as digoxin and myosin activators such as MYK-491 could lead to an elevated mass and decreased eccentricity. Experimental studies on hypertrophic cardiomyopathy indicated Ca^2+^ channel blockers like diltiazem^56^ and myosin inhibitors like mavacamten^69^ could reduce cardiac mass and increase cardiomyocyte eccentricity based on measured chamber dimensions. The angiotensin receptor blockers such as losartan^70^ and candesartan^71^ also could decrease cardiac mass, but their effect on cardiomyocyte eccentricity was non-significant.

**Table 1.**
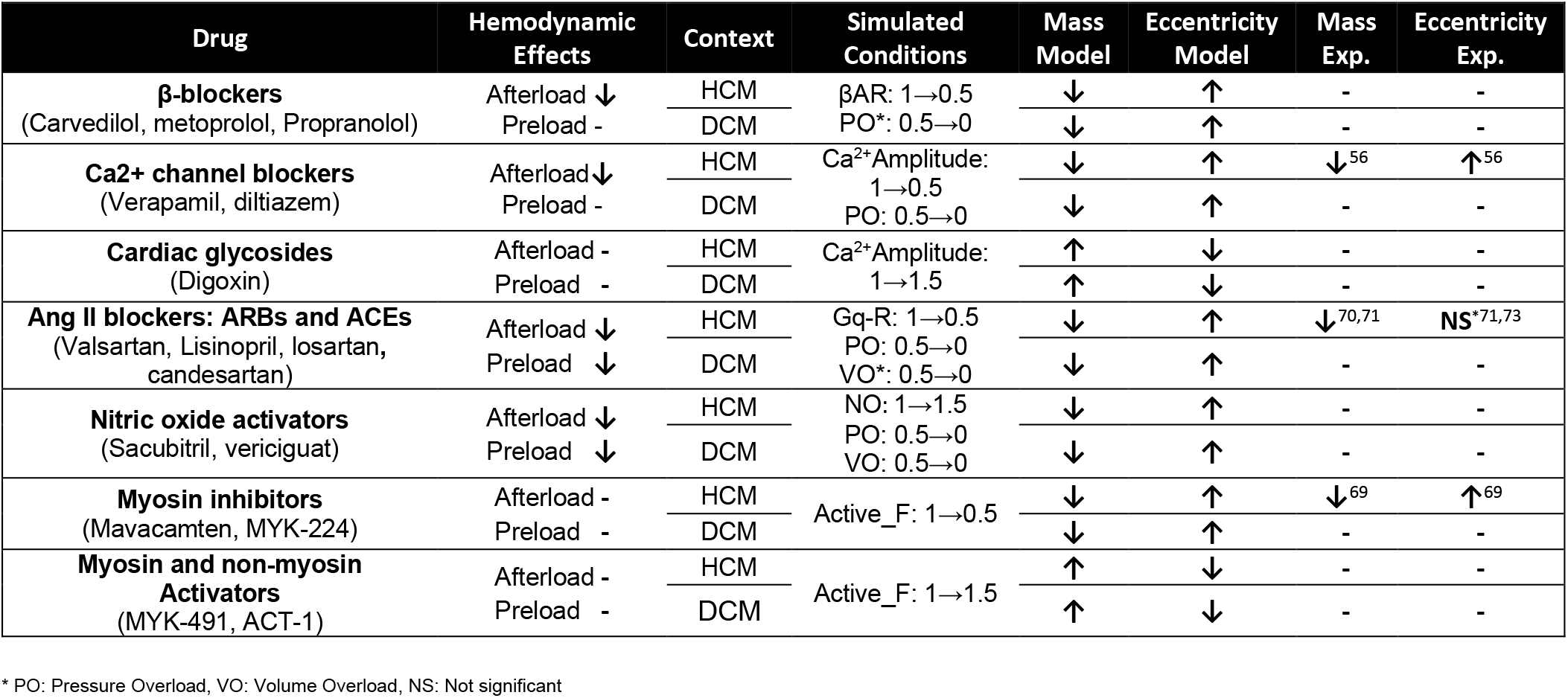
Potential effects of existing drugs on cardiomyocyte morphology in HCM/DCM contexts.

Drug-induced hemodynamic changes could be a significant part of drugs’ effects on cardiomyocytes morphology and reverse cardiac remodeling in heart failure^72^. To explore this in familial cardiomyopathy, we compared the effects of medications from Table 1 on cardiomyocyte morphology with and without considering their hemodynamic changes. As shown in supplementary Table S3, excluding the altered hemodynamics of drugs does not change their effects on cardiomyocyte morphology qualitatively. However, depending on the drug, it could significantly reduce the strength of the drug’s influence on cardiomyocyte morphology. For example, as shown in S3 Table, excluding the hemodynamic effects of nitric oxide activators leads to non-significant changes in cardiomyocyte morphology due to nitric oxide activators.

Moreover, it is noteworthy that Table 1 only includes altered features that the model was designed to simulate. Thus, drug responses like altered cardiac workload that are regulated by complex mechanisms including endocrine and paracrine signaling pathways, autonomic nervous system, and metabolic pathways^74,75^ are not covered in Table 1. However, the current model covers the main signaling pathways in familial cardiomyopathy as well as the effects of pressure and volume overload stimuli.

## 3. Discussion

Here, we proposed a conceptual model to explain distinct phenotypes in familial cardiomyopathy. Inspired by previous experimental studies, we illustrated in the conceptual model how cardiac active and passive forces and titin stiffness are linked to myocyte eccentricity in different conditions. By considering the central role of titin and its stiffness in regulating cardiac response in familial cardiomyopathy, we developed and validated a predictive model of the HCM/DCM signaling network to understand the underlying mechanisms of genotype to phenotype in familial cardiomyopathy. The model sensitivity analysis revealed influential reactions regulating the HCM/DCM signaling network and suggested new potential interactions that increase model prediction accuracy. We performed a combination pharmacotherapy analysis through *in-silico* knockdown and overexpression of signaling nodes. We discovered potential drug targets to inhibit cardiac growth and morphology changes in familial HCM and DCM. We experimentally validated a potential drug combination predicted by the model (inhibition of ERK+PI3K-AKT) with patient-specific iPSC-CMs, showing its effectiveness in reversing HCM phenotypes. We also simulated the effects of current medications on cardiomyocyte eccentricity and mass and evaluated their effectiveness. This HCM/DCM signaling model provides a framework to investigate genotype to phenotype mechanisms in familial cardiomyopathy. The model yields insights into new potential drugs to decrease the maladaptive cardiac growth and remodeling in familial cardiomyopathy as the main precursor of heart failure.

### 3.1 Titin and cardiomyopathy

In addition to the active force module, the model analysis indicated titin could play a significant role in regulating HCM/DCM phenotypes. Titin is considered an important mediator in initiating and developing familial cardiomyopathy^76^. However, underlying mechanisms by which titin links genotype to phenotype in familial cardiomyopathy are not fully understood. As shown in the model (Fig. 1b), kinases such as PKD, PKG, CaMK, PKA, and ERK1/2 phosphorylate titin and reduce its stiffness, contrary to PKC that increases titin stiffness by phosphorylation^77^. In HCM, while the model predicted an increase in the activity of PKD, PKG, CaMK, and ERK1/2 kinases consistent with experimental data^15,27,29^, both model and experimental studies showed no decrease in titin stiffness^29,78^. This highlights the importance of other factors such as elevated PKC or alteration in titin splicing as causes of the long-term increase in titin stiffness during HCM. In DCM, the model predicted a decrease in the activity of all kinases except PKA. However, experimental studies depicted discordant kinases activity in ERK1/2^11,21^ and CaMK activity in DCM^21,37^. These could be the results of diversity in DCM gene mutations^6^, transient response of ERK1/2 activity, and activation of CaMK by other pathways^79^. The projected decrease in titin stiffness by model and patients with dilated cardiomyopathy^42^ suggests the significance of PKC-dependent phosphorylation of titin’s PEVK domain^77^ and other non-kinase regulators in altering titin stiffness in DCM.

RBM20 is a key player in controlling the splicing process of titin mRNA^80^, and RBM20 mutation results in DCM. Inhibiting RBM20 function leads to an increase in the more compliant N2BA titin isoform and decreases titin stiffness^81^. Rbm20ΔRRM heterozygous mice have higher LV chamber compliance and enhanced exercise performance with normal systolic function.^81^ As shown in the HCM/DCM signaling model (Fig. 1b), mTOR could link PI3K-AKT and ERK1/2 pathways to RBM20. The model indicates an increase in mTOR activity in all contexts, supported by an Ikeda et al. study of volume overload^82^ and a study by Yano et al. on DCM^83^. Elevated mTOR activity could increase titin stiffness through RBM20 as observed in all contexts except DCM^42^. In DCM, decreased titin stiffness could be due to the high expression of compliant N2BA isoform^42,84^, which suggests the role of other regulators of the titin splicing process or stiffness in DCM patients. As shown in the model, calcium can modify titin stiffness, too. Recent studies displayed elevated titin force and stiffness in calcium-activated myofibrils^85^. While calcium alone raises titin stiffness^86^, its influence is small compared to increased titin stiffness after sarcomere activation^87^. Nishikawa et al.^85^ described that titin–thin filament interaction in active sarcomeres is mainly responsible for this calcium-dependent increase in titin stiffness. Thus, part of the change in titin stiffness, especially in DCM, could be due to altered calcium predicted by the model.

Furthermore, titin is a signal transducer in cardiac mechanotransduction^88^. Among the titin-associated proteins, FHL1 and FHL2 are decisive in HCM/DCM signaling and activate ERK1/2 and NFAT pathways. In agreement with experimental data (Fig. 2), the model illustrated an increase in FHL1 activity in HCM, pressure overload, and volume overload contexts, which then could increase ERK1/2 activity and regulate cardiomyocyte eccentricity. Regarding FHL2, HCM mutation decreases FHL2 mRNA and protein levels^89^ and thus attenuates its inhibitory effect on NFAT through binding to CaN^90^. Moreover, the model considers titin’s contribution to active force development. Modifying the Frank-Starling mechanism by altering interfilament lattice spacing^91^, titin stiffening in muscle activation, and unfolding and refolding titin IgG domains^92^ are potential mechanisms that could link titin to HCM and DCM phenotypes.

### 3.2 CaMK: Integrator of calcium signaling in familial cardiomyopathy

The multifunctional signaling molecule CaMK has been acknowledged as a master regulator of arrhythmias and maladaptive remodeling in cardiac diseases^93^. As predicted by the model (Fig. 3), CaMK acts as an integrator of altered calcium signals and regulates genotype to phenotype mechanisms in cardiomyopathy. CaMK completes a control loop in calcium signaling by linking the changes in calcium transient to the activation of calcium cycling regulators such as SERCA, LCC, and RyR. Moreover, it alters cardiomyocyte stiffness and calcium sensitivity through titin and troponin phosphorylation, respectively, and can affect cardiomyocyte morphology through ERK1/2 and MEF2^94^. Recently, various studies have presented CaMKIIδ as a target of RBM20 that might be involved in RBM20 mutation-induced dilated cardiomyopathy^95^. While some studies reported the beneficial effects of CaMK modification^96^, altering CaMK activity as an effective treatment strategy in cardiomyopathy (Fig. 5) or even heart failure^97^ is challenging due to a large number of CaMK target proteins. Though a consistent increase in CaMK activity has been reported for the HCM context^27,98^, reported CaMK activity in the DCM context is variant^21,38^. This inconsistency may arise from diverse gene mutations leading to DCM that initiate different compensatory mechanisms to regulate calcium cycling and the level of disease progression. Robinson et al.^21^ studied functional consequences of three variant mutations in thin-filament regulatory proteins that led to DCM with reduced systolic Ca^2+^, SERCA, and CaMK activity. However, Liu et al.^38^ reported decreased SERCA and Ca^2+^ transients with increased CaMK activity after R25C-PLN mutation in the DCM.

Studies investigating DCM in patients often do not differentiate between various etiologies. Furthermore, data from patients with end-stage heart failure makes it harder to find the primary reason for the reported increase in CaMK activity^99^. To sum up, CaMK could be a key drug target in familial cardiomyopathy due to its central role in HCM/DCM signaling (Fig. 3). However, a wide range of mechanisms affecting CaMK activity in human dilated cardiomyopathy and heart failure^94^ and contradictory impacts of CaMK on different mediators of HCM/DCM signaling demands more specific studies on CaMK to predict its therapeutic effects accurately.

### 3.3 Challenges in drug development

The main challenges in developing drugs for familial cardiomyopathy are the significant diversity in the genotype to phenotype relationship and its unknown mechanisms. This study explored part of these mechanisms concerning the cardiac signaling network and predicted some potential drug targets to decrease cardiac growth and remodeling in familial cardiomyopathy. According to the model predictions, inhibition of PI3K-AKT or NFAT seem the most effective target for inhibition of elevated cardiac mass during HCM and DCM. In this regard, Fan et al.^100^, in a murine model study, showed that DM-celecoxib and Celecoxib, which inhibit AKT and then NFAT, improved left ventricular systolic functions and prevented cardiac remodeling and reduced mortality in DCM. However, considering no available clinical study to support these observations and the existence of other reports showing the opposite effects of AKT knockout in murine models leading to pathological hypertrophy^101^, more studies are needed to explore the impact of AKT and NFAT modifications on familial cardiomyopathy.

Moreover, the efficacy of inhibiting PI3K-AKT and GF-Rs has been established in blunting hypertrophic response to exercise training^102^, pressure overload^103^, and neurohormonal factors^104,105^. The role of PI3K-AKT and EGFR inhibition in preventing cardiac fibrosis, a significant characteristic of HCM, has also been reported. The main concern in inhibiting GF-Rs, especially PI3K-AKT, is their vital role in other cardiac functions such as cardiomyocyte proliferation, survival, and apoptosis^106^. mTOR is another potential drug target for HCM (Fig. 5). Rapamycin, a specific mTOR inhibitor, could attenuate or reverse cardiac growth and remodeling in pressure overload^107^ and HCM^108^ contexts. Altering titin stiffness is another effective treatment strategy in HCM and DCM, as predicted by the model (Fig. 5). While HCM mutations led to a higher stiffness in both titin and cardiac tissue (ECM), the model results suggest that more compliant titin could partly reduce hypertrophic remodeling and subsequently improve its diastolic dysfunction. Methawasin et al.^109^ showed a significant improvement in diastolic dysfunction and exercise tolerance after reducing titin stiffness in pressure-overload hypertrophy leading to HFpEF. In DCM, model analysis suggests that an increase in titin stiffness could be beneficial by lowering cell eccentricity. Hutchinson et al.^110^ showed that increased myocardial stiffness in the volume overload context is beneficial and limits its eccentric remodeling.

### 3.4 Limitations and future directions

Although the model effectively predicts the variations of signaling pathways that lead to familial HCM and DCM, it has some limitations. The cardiac metabolic network plays a significant role in mediating genotype to phenotype in familial cardiomyopathies, especially in their progression to heart failure^111^. In DCM, truncating mutations in the TTN gene were associated with a reduced rate of O2 consumption, significant ROS production, and transcriptional upregulation of mitochondrial oxidative phosphorylation system^112,113^. Galata et al.^114^ reported an association between altered ERK1/2 signaling after LMNA gene mutation and changes in shape, fragmentation, and distribution of cardiomyocyte mitochondria. In HCM, deficient creatine kinase-phosphocreatine coupling in the hypercontractile myocardium led to diastolic dysfunction through inadequate ATP regeneration^115^.

Furthermore, as shown in Table 1, the model results indicate reduced mass and increase in the eccentricity of cardiomyocytes after applying the effects of some heart failure drugs such as beta-blockers, ACE inhibitors, or ARBs. These results are not consistent with several experimental findings, especially in patients with idiopathic DCM that illustrate the improvement of cardiac left ventricle morphology and function (reverse remodeling) in some patients^116,117^. As the current model was developed to investigate the role of cardiomyocyte signaling in linking mutations-induced changes in calcium sensitivity (primary factor) to cardiomyocyte morphology, there are limitations in extending model results to clinical and tissue scale data. Clinically relevant aspects of cardiomyopathy such as systolic or diastolic dysfunction, fibrosis, and apoptosis contribute to heart morphological changes in tissue level and affect heart response to a specific medication. These changes could also modify the biomechanical forces sensed by cardiomyocytes and affect phenotype development^118^. In addition to model limitations, significant variations in familial cardiomyopathy mutations and the variable clinical effectiveness of drugs should be considered when comparing model results with clinical data. The most available literature on clinical data for patients with idiopathic DCM does not report the cause of DCM. In this regard, Hershberger et al.^6^ stated that familial DCM could be identified in only 20-35% of patients with idiopathic DCM. Thus, interpretation of data and effectiveness of drugs in familial cardiomyopathy, especially on idiopathic DCM, should be done with care.

In addition, signaling pathways regulating the cardiomyocyte morphology during familial cardiomyopathy are generally dynamic and act through different time courses in the process of cardiomyocyte growth and remodeling. As rich kinetic quantitative data are not available for these signaling pathways during familial cardiomyopathy, the model does not consider these temporal responses. However, despite this limitation, the model validates remarkably well (see Fig. 2) as this model was developed based on causal interactions between signaling components in cardiomyocytes and uses default parameters instead of a specific dataset solving the problem of overfitting to particular stimuli and timing. Also, since all molecular mechanisms connecting sarcomere force generation to signaling pathways in familial cardiomyopathy are not discovered yet and other cellular mechanisms like metabolic pathways shown to be significant in the determination of primary versus secondary signals in familial cardiomyopathy, the current model does not predict primary versus secondary signals. However, in our model change in active force due to altered calcium sensitivity is considered as the primary signal. On the condition of more quantitative data on different scales of molecular mechanisms regulating cardiomyocyte response in familial cardiomyopathy, a multi-scale model could address these limitations and predict the influence of mutation-induced changes in sarcomere dynamics on signaling pathways by considering the role of more complex phenomena such as myosin super relaxed to a disordered relaxed state change^119^ in familial cardiomyopathy.

For the future, integrating the HCM/DCM signaling model with a cardiac metabolic network as well as incorporating variations in the cell mechanics and mechanotransduction in the model would be beneficial. Additional in-vitro experiments on human iPSC-derived cardiomyocytes to test the effects of other potential drug targets suggested by the model could confirm the accuracy of the model predictions. Finally, we developed and validated the HCM/DCM signaling model, which provides new insights into the genotype-to-phenotype mechanisms in familial cardiomyopathy.

## 4. Methods

### 4.1 Model construction

A knowledge-based computational model of the HCM/DCM signaling network was developed from the literature. Main cardiac signaling modules were identified from over 40 peer-reviewed papers addressing the role of certain signaling modules in dilated or hypertrophic cardiomyopathy by *in-vitro* and *in-vivo* studies. Individual reactions between signaling nodes were then added, based on cardiac signaling networks developed by Ryall et al.^24^ and Tan et al.^25^ that modeled cardiac hypertrophy and mechano-signaling networks, respectively. In total, 124 references were employed for model development. These references were not used for the model validation. In the calcium module, we focused on steady-state variations in the activity of its major regulators such as RyR, SERCA, PLB, LCC and their effects on Ca^2+^ transient characteristics such as Ca^2+^ amplitude, diastolic level, decay time, and Ca^2+^ buffering and SR Load. Calcium average and active force nodes are integrated parameters to consider the time and amplitude of their variations^11^.

We employed the logic-based differential equation (LDE) approach^37^ to build a predictive framework to explore HCM/DCM signaling dynamics. A normalized Hill function modeled activation of each node by its upstream reactions. We modeled pathways crosstalk by continuous gates representing “OR” and “AND” logic^37^. The OR gates were used for reactions that modify the node regardless of others and the AND gates for reactions affecting each other. The reaction parameters in the model are reaction weight (W=0.5), half-maximal effective concentration (EC_50_=0.5), and Hill coefficient (n=1.4). The time constant (τ=1), initial activation (Yinit=0), and maximal activation (Ymax=1) regulate the dynamics of signaling nodes. In the case of multiple inhibitory inputs for a node, such as titin stiffness or calcium sensitivity, the default reaction weight leads to an oversaturated node activity.

Furthermore, a default weight for the reaction directly linking DCM-Mutation to PI3K-AKT activation makes PI3K-AKT insensitive to its other upstream regulators. Thus, we used a reaction weight of 0.1 for these reactions (S1 Table). The Netflux software (available at https://github.com/saucermanlab/Netflux) has been used to generate a system of LDEs. Simulation of experimental conditions has been performed by the model validation module (available at https://github.com/mkm1712/Automated_Validation) in MATLAB software.

### 4.2 Context-specific data collection

Experimental data of familial hypertrophic and dilated cardiomyopathies, pressure overload, and volume overload conditions were manually curated from the literature. We acquired qualitative data at steady-state or appropriate time points for nodes with transient activity from over 70 research articles with similar cell lines and experimental conditions. Overall, 160 qualitative experimental data sets with and without overexpression and inhibition perturbations were obtained, including 41 and 48 data points for HCM and DCM contexts. The context-specific data, simulated conditions, and references are displayed in the S2 Table.

### 4.3 Global sensitivity analysis

We employed a Morris global sensitivity analysis approach to identify signaling reactions with a greater influence on the model validity in familial cardiomyopathy. The Morris elementary effects method is a statistical method to screen models with many parameters or high computational costs for simulation. We utilized the Sampling for Uniformity (SU) strategy with input factor level equal to eight, oversampling size of 300, and trajectories’ number equal to 16 to generate the sampling data using the EE sensitivity package developed by Khare et al.^120^. The reaction weight (W) was selected as the variable parameter to compute the sensitivity measure, μ*, for each reaction. μ* is the average of the elementary effects’ absolute value to certify robustness against non-monotonic models but cannot capture nonlinear effects. Therefore, to account for interactions between factors or nonlinearities in the model, we computed σ as the standard deviation of the elementary effects.

### 4.4 Human iPSCs

Detailed information of human iPSCs were reported^65^. Briefly, under Yale University Institutional Review Board’s approval, MLP-W4R;MYH7-R723C iPSC lines were generated by infecting peripheral blood mononuclear cells (PBMC) with Sendai virus particles encoding human transcription factors of SOX2, OCT4, KLF4, and c-MYC. The iPSCs line were cultured on the mouse embryonic fibroblast feeder (MEF). After reaching 70-80% confluency (~4-5 day), cells were dissociated from culture dishes utilizing 1 mg/mL of Dispase II (Gibco; 17105041) and spun down at 200 rpm for 4 minutes. iPSCs were then replated on Growth Factor Reduced (GFR) Matrigel-coated (Corning; 1:60 diluted in DMEM/F12) plates. Cardiac differentiation was started when iPSCs reached 80-95% confluency (in about ~1-2 days). Cardiac differentiation was conducted as previously described^121^.

### 4.5 Drug treatments and analysis

On Day 25, iPSC-CMs were treated with 0.5 μM MEK-ERK inhibitor (U0126-EtOH, Selleck; S1102), 0.2 μM PI3K-AKT inhibitor (Wortmannin, Selleck; S2758), or DMSO vehicle control. The drug doses were determined to minimize iPSC-CMs toxicity. After 4 days drug treatment (day 29) cardiomyocytes were fixed with 4% paraformaldehyde (ProSciTech; EMS15714), washed with Dulbeccos Phosphate Buffered Saline (DPBS, Sigma-Aldrich; D8537), and permeabilized with 0.1% Triton X100 (AmericanBio, Inc; 9002-93-1). iPSC-CMs were then stained by mouse anticTnT (ThermoFisher Scientific; MS-295-P0). Imaging was conducted by an inverted fluorescent microscope (Olympus; CKX53). Cell area was measured by ImageJ from a minimum of 3 independent batches of cardiac differentiation.

For messenger RNA (mRNA) analysis, after 24 hours drug treatment, total RNA was isolated from iPSC-CMs using TRIzolTM kit (ThermoFisher Scientific; 12183555). The cDNA synthesis was performed by employing iScriptTM cDNA synthesis kit (Bio-Rad Laboratories; 1708891) following manufacturer’s protocols. Quantitative PCR was conducted using IQTM SYBR Green Supermix (Bio-Rad Laboratories; 1708882) on CFX96 Optical Reaction Module for RealTime System (Bio-Rad Laboratories; 1845096). All experiments were conducted from a minimum of 3 independent batches. Real-time sequence detection software (Bio-Rad Laboratories; 1845001) was used to compute Cycle threshold (CT). Analysis was performed using ΔΔCT method^122^ and fold change was computed.

## Acknowledgments

The authors thank Dr. Jennifer Davis for her valuable feedback. This study was funded by the awards received by the National Institutes of Health (HL137755 to J.J.S.; HL137100 to A.D.M.; HL164783 to Y.Q.) and the University of Virginia Pinn Award to J.J.S. The funder had no role in study design, data collection, analysis, decision to publish, or manuscript preparation.

