## Supplemental Table 3 for "Signaling network model of cardiomyocyte morphological changes in familial cardiomyopathy"

**Table 1 Evaluation of pharmacotherapies predicted by the model**

| Context | Perturbation (drug) | Targeted Component (s) | Model Prediction (Fig. 5) | Experimental  Outcome |
| --- | --- | --- | --- | --- |
| DCM^1^ | Overexpressed MEK1 | Activation of ERK1/2 | ✓ | ✓ |
| HCM^2^ | PD0325901 | Inhibition of ERK1/2 | ✓ | ✓ |
| HCM^2^ | Rapamycin | Inhibition of mTOR | ✓ | ✓ |
| HCM^2^ | Carvedilol | Inhibition of BAR (& Calcium) | ✓ | ✓ |
| 1 Davis et al.^11^  2 Bu et al.^55^ | | | | |
